# Analysis of multi-trait evolution across independently evolved cavefish populations reveals shared and independent evolution of suites of cave-associated traits

**DOI:** 10.1101/2025.10.22.683983

**Authors:** Stefan Choy, Maya Enriquez, Rianna Ambosie, Aubrey E. Manning, Briley F. Mullin, Roberto Rodriguez-Morales, Jennah Abdelaziz, Sarah Jacobson, Evan Lloyd, Alli Kimmel, Solomia Lapko, Isabel Carino-Bazan, Helena Bilandžija, Alex C. Keene, Erik Duboue, Suzanne McGaugh, Johanna E. Kowalko

## Abstract

Environmental perturbations often lead to the evolution of multiple traits. Determining whether shared genetic factors underlie multi-trait evolution is a central question in evolutionary biology. In the Mexican tetra, *Astyanax mexicanus,* cave-dwelling populations have repeatedly evolved multiple traits. The repeated evolution of these traits, paired the robust environmental differences between the surface and cave habitats, provide an opportunity to investigate the genetic basis of multi-trait evolution. Here, we investigate the extent to which shared genetic mechanisms underlie the repeated evolution of multiple traits in cavefish. Across cave populations, we find evidence for shared and distinct genetic mechanisms contributing to the evolution of individual traits. Further, multiple traits covary in cave-surface F2 hybrids and many of the same trait correlations are found across independently evolved cave populations. Finally, we assessed traits that differ between pigmented and albino F2 fish in surface fish with mutations in the albinism gene *oculocutaneous albinism 2 (oca2)*. This revealed that mutations in *oca2* reduce bottom-dwelling behavior in *A. mexicanus*. Together, these findings suggest that multi-trait evolution occurs repeatedly through shared genetic factors across *A. mexicanus* cave populations. These results are consistent with pleiotropy or linkage playing a large role in multi-trait evolution in this species.

## Introduction

Populations inhabiting similar environments or under similar environmental pressures frequently evolve similar phenotypes. For example, in two species of pocket mice and a species of fence lizards, divergent coloration has evolved between different populations that live on different colored substrates such that individuals’ colors matches their substrate, reducing predation [1]. While the environmental pressures leading to repeated evolution have been studied extensively, the extent to which shared genetic mechanisms drive repeated evolution is less understood.

Even among similar traits, examples of both shared and distinct genetic mechanisms contributing to repeated evolution have been observed. Resistance to the poison in milkweed has independently evolved in at least three species, including the monarch butterfly, the red milkweed beetle, and oleander aphids, through mutations in the same gene encoding NA,K-ATPase [2,3]. In contrast, yellow pigmentation in *Drosophila novamexicana* and *D. americana* evolved via distinct genetic mechanisms [4]. While traits may evolve through shared or distinct genetic mechanisms, more research is needed to understand why traits evolve through shared mechanisms in some instances, but independent mechanisms in others.

One factor that may influence how frequently a locus is utilized during evolution is pleiotropy. Historically, pleiotropic loci — loci that impact multiple traits — have been considered maladaptive in evolutionary contexts [5–7]. However, recent work suggests that pleiotropic loci are beneficial under certain conditions, such as when there is maladaptive gene flow or if a pleiotropic locus results in alterations to a suite of traits that together have positive fitness consequences [8–10]. Indeed, pleiotropic loci can drive the evolution of multiple traits. For example, when invading freshwater environments, sticklebacks evolved not only freshwater tolerance, but also reduced plate armor, body position when schooling, and lateral line patterning through the pleiotropic *edar* gene [11–16]. Likewise, close linkage can promote multi-trait evolution in cases where independent genes produce adaptive changes in multiple traits [8,17]. Thus, multiple traits may evolve through shared genetic loci repeatedly due to the benefits of linkage and/or pleiotropy. Here, we directly test the extent to which repeated evolution of multiple traits occurs through the same genetic loci in the blind cavefish *Astyanax mexicanus*.

The freshwater fish *Astyanax mexicanus* is a powerful system for investigating repeated evolution. *Astyanax mexicanus* exists as two primary morphotypes: an eyed, riverine surface fish and a blind, subterranean cavefish. At least 35 distinct populations of cavefish have been identified, which derive from at least two colonization events from surface ancestors [18–20]. Cave populations have repeatedly evolved multiple behavioral, physiological and morphological trait changes relative to surface fish, including reduced sleep, reduced stress response behaviors, eye loss, albinism, and increased numbers of superficial neuromasts, sensory organs of the lateral line [12–19]. Additionally, cavefish remain interfertile with surface fish, enabling the generation of hybrids to investigate the genetic basis of specialized cave traits [20]. Together, these advantages make *A. mexicanus* a powerful model for investigating multi-trait evolution in response to the repeated colonization of a new environment.

Both genetic mapping studies and analysis of covariance of multiple cave-evolved traits in cave-surface hybrid fish suggest that concentrated genetic architecture may contribute to the evolution of cave traits in this species [21–28]. For example, quantitative trait loci (QTL) for different traits overlap in the *A. mexicanus* genome, and this occurs more frequently than expected by chance [21–24]. Moreover, functional studies demonstrate that the gene underlying albinism in *A. mexicanus* cave populations, *oculocutaneous albinism 2* (*oca2*), likely impacts at least four other cave-evolved traits, catecholaminergic signaling, anesthesia resistance, reduced sleep and larval prey capture behavior [27,29–31]. In addition, the gene *sonic hedgehog* has been implicated in the evolution of multiple cave traits, including eye degeneration, expansion of the jaw and tastebuds, and alterations to the forebrain [32–34]. Together, these studies suggest that pleiotropy or linkage may be significant drivers of adaptation to cave life in *A. mexicanus* cavefish. However, most prior genetic studies have focused on only one cave population at a time. Thus, whether evolution of multiple traits through shared genetic architecture has occurred repeatedly across *A. mexicanus* cave populations is currently unknown.

To systematically investigate the relationship between the genetic architecture of cave-derived traits, we established a pipeline to phenotype nine traits in surface fish, cavefish, and surface-cave and cave-cave hybrid populations. We assessed trait correlations in hybrid populations to determine whether multiple traits evolved through concentrated genetic architecture repeatedly across cavefish evolution. Further, we investigated covariations between albinism and other traits directly by assessing bottom dwelling and eye size in *oca2* mutant surface fish. Together, this work suggests that shared genetic architecture between discrete traits may be a central driver of the repeated evolution of the cave phenotypes in *A. mexicanus*.

## Methods

### Fish husbandry

Husbandry of *A. mexicanus* was carried out based on established methods [44,45]. Briefly, larvae were housed in incubators at 25°C on a 14:10 light:dark (L/D) schedule in glass bowls until 6 days post fertilization (dpf). Starting at 6dpf, *A. mexicanus* larvae were moved to plastic tanks filled with 2 liters of water at a density of N = 30 per tank and were fed brine shrimp twice a day. Between 14-30dpf, larvae were moved into 2.8L plastic tanks on an Aquaneering recirculating water system. After 30dpf, larvae were fed Gemma Micro 300 food (Skretting) in addition to brine shrimp. Larvae were size-sorted every 2-4 weeks. After reaching approximately 3cm in length, fish were transferred into 5-gallon tanks where they were fed 1mm Ziegler pellets twice a day and kept at a maximum density of 15 fish per tank. Water temperature was maintained at 23 ± 1°C. Prior to behavioral assays, fish were fed one time per day for at least two weeks.

### Hybrid generation

A nested hybrid population was generated by crossing individual *A. mexicanus* males from the Pachón, Molino, and Tinaja cave populations to the same female surface fish. This crossing structure generated surface-Pachón, surface-Molino, and surface-Tinaja F1 hybrids that were all half-siblings. F2 hybrids were generated by incrossing F1 hybrids within each F1 cross (i.e., not between crosses), where all F2 hybrids share the same F0 surface fish ancestor.

### Phenotyping pipeline

Surface fish, Pachón, Molino, and Tinaja cavefish, hybrid fish were assayed for nine behavioral, physiological, and morphological traits (Fig 1A). These traits were stress response via a novel tank test to assess bottom-dwelling, sleep over 24 hours, weight loss after 30 days of starvation, sleep over 24 hours after 30 days of starvation, anesthesia resistance, eye size, pupil size, cranial superficial neuromast number, and the presence or absence of pigmentation (Fig 1). A subset of individuals went through a shortened pipeline where the 30 day-starvation period was omitted. These individuals do not have measurements for day 30 sleep or weight loss but were assayed for anesthesia resistance and morphological traits following day 0 sleep. For a separate project, fish were assayed for aggression via resident-intruder assays the day following novel tank assays and recorded for 24 hours to ascertain sleep phenotypes at days 0, 15, and 30 of starvation; only the days 0 and 30 sleep data are reported here.

**Figure 1.**
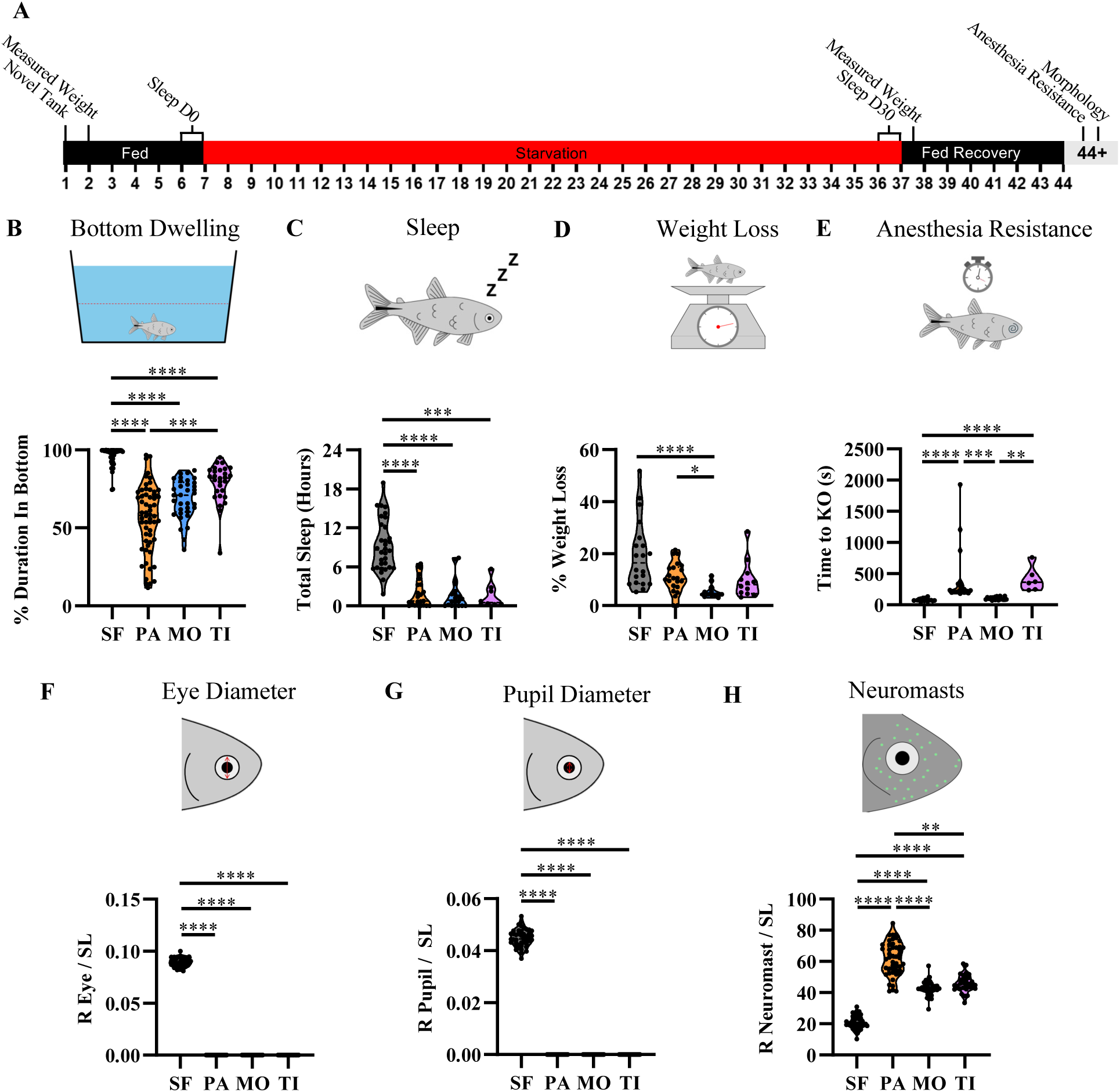
Cave populations of *A. mexicanus* repeatedly evolve traits that differ from surface populations. A) Overview of the experimental pipeline that was used to assay surface fish, cavefish, and hybrids. Eye morphology, neuromasts, and standard length data was captured during the ‘morphology’ section. B) Percent time spent bottom-dwelling during the novel tank. C) The percent weight lost after 30 days of starvation. D) Time spent awake in the presence of tricaine anesthetic E) Dorsal-ventral right eye diameter corrected by standard length. F) Dorsal-ventral right pupil diameter corrected by standard length. G) Number of superficial neuromasts located overtop of the right suborbital 3 bone corrected by standard length. Levels of significance are represented with asterisks: p < 0.05 = *, p < 0.01 = **, p < 0.001 = ***, p < 0.0001 = ****.

### Novel tank

Bottom-dwelling in a novel tank was assessed as a measure of stress, as previously described [27]. Fish were removed from their home tank and placed in a 500ml beaker to acclimate for 10 minutes. The water and fish were then gently transferred to a new tank (21.3cm x 12.7cm x12.7cm) filled with 2 liters of water. Warm white LED strip lights were used to illuminate the novel tank setup. The 10 minutes following fish placement into the assay tank were recorded at 30 frames per second (fps) with a Basler ace acA1300-60gm GigE Mono camera (Edmund Optics: 88327) with Basler PylonViewer software. The novel tank videos were analyzed via Ethovision (version 17.0.1630), an automatic tracking software. In Ethovision, the analysis arena was bisected into top and bottom zones. Time spent in the bottom zone was converted to a percent of the total and reported. Videos for which the entire 10-minute recording did not complete, or for which Ethovision was unable to track the fish for more than 10% of the total video were excluded from further analysis.

### Sleep and starvation

Following the novel tank assay, fish were weighed to ascertain pre-starvation weight and moved into experimental sleep tanks (45 x 17 x 17cm) [46]. Tanks were divided into five segments using opaque dividers and one fish was placed into each of the first four segment, while the fifth segment housed a sponge bubbler. All assays were backlit with 850nm infrared strip lights and recorded with a modified Microsoft LifeCams (Amazon: B004ABO7QI) fitted with an IR-pass filter, so that illumination was constant between light and dark conditions. Overhead warm white LED lights were set on a 14:10 L/D light cycle. Fish were allowed to acclimate for four days and were tested on the fifth day of being in the tank for initial sleep phenotyping. Fish were fed bloodworms once per day during acclimation. Fish were then starved for 30 days, during which water changes with conditioned water were performed every 5 days. After the 30-day starvation period, fish spent another 7 days in the segmented tanks and were fed bloodworms once per day. A subset of parental populations (surface fish, Pachón, Molino, and Tinaja) were fed bloodworms once per day to serve as controls.

The recorded sleep videos were processed using Ethovision behavior tracking software to quantify the amount of time spent sleeping based on velocity measurements [47,48]. The utf-16 encoded data files were then converted into utf-8 files and processed via a custom python code (see Supplemental Files) to extract the amount of time spent sleeping. Specifically, fish found to be moving at less than 4 cm/s for more than 60 or more seconds were considered to be asleep, as in [47]. Only 23.5 hours were analyzed, with analysis starting 30 minutes after lights on (zeitgeber time 0.5). Here, we report the total hours of sleep across the 23.5 hours.

At the beginning of acclimation for sleep, the end of starvation, 7 days post starvation (after recovery), and immediately before anesthesia resistance, fish were weighed by placing individual fish into a water-filled beaker on a top-loading balance. Fish weights were recorded in grams up to one significant figure. 40 fish demonstrated a weight gain during the 30 days of starvation, and these data were removed under the assumption that a measuring error was made.

### Anesthesia resistance

Response to anesthesia was measured after at least seven days of recovery from starvation, as previously described [39]. Fish were placed into a 500mL beaker filled with 200mL conditioned water and allowed to acclimate for at least 10 minutes. Following acclimation, 200mL of 200mg/L MS-222, or tricaine, conditioned water solution, adjusted to proper pH with sodium bicarbonate, was added to the beaker, and the time to unconsciousness was recorded to the nearest second. Unconsciousness was defined as the fish no longer responding to prodding with a plastic pipette to the head and operculum. Following these trials, fish were immediately returned to their home tank and allowed to recover.

### Morphology

Fish were stained with a 28.9mg/L DASPEI solution for 60 minutes in the dark to prevent breakdown of DASPEI, as previously described [49]. Following a brief treatment with chilled tricaine, photos of the lateral view of the head and fluorescently labelled neuromasts were taken on a fluorescent dissecting microscope (Leica M165 FC). Additionally, full-body photos were taken with a Canon Rebel T7 camera (Amazon: B07C2Z21X5) with a centimeter ruler and color standard in frame for standard length quantification. Standard length was measured from the tip of the snout to the base of the tail using FIJI [50].

### Eye analysis

Eye and pupil sizes were quantified by measuring the dorsal-ventral diameter of the eye and pupil using FIJI [50]. A subset of eyes or pupils from F2 hybrid fish were not fully visible (obscured by skin, occluded by orbital bone, etc.) and were not quantified, as the edges of the structure were not identifiable. Both eye and pupil size were correlated with fish length, thus, both metrics are reported as corrected for standard length by dividing by standard length.

### Neuromast analysis

Superficial neuromasts within the boundary of the third suborbital bone (SO3) were quantified on each side of the fish via a custom ImageJ code [51] (see Supplemental Materials). The quantification of neuromasts was limited to the SO3 area since neuromasts in this section of the animal have previously been shown to directly correlated with food finding abilities and sleep [31,48]. The code has users outline the SO3 anatomical area and then utilizes a background subtraction algorithm to count neuromasts within this boundary. Each image was then manually checked and corrected to ensure accurate neuromast quantification. Any images that exhibited signs of DASPEI overstaining or fish that had anatomical abnormalities that significantly warped the SO3 area were excluded from analysis. The anatomical right standardized neuromasts were used in this analysis, with the left side being reported in the supplemental figures. As the number of neuromasts correlated with the size of the fish, we reported corrected neuromast number (number divided by standard length) for this trait. However, we also reported values for total neuromast number in the supplemental figures.

### oca2 mutant surface fish

To directly test the role of pleiotropy in multi-trait evolution of *Astyanax* cavefish, we used a line of surface fish with a CRISPR-Cas9-induced 2 base pair (bp) deletion in exon 21 of *oca2* [36,52]. Here, fish heterozygous for the 2bp deletion were incrossed, and offspring were raised to adult stages (>6 months). Albino and pigmented fish were raised separately to prevent competition. All offspring from incrosses were assayed in the novel tank assay and then imaged using a Canon Rebel camera. After data collection, fish were anesthetized, and part of the tail fin was clipped for DNA extraction to genotype the pigmented individuals to distinguish heterozygous individuals from wild-type individuals [36,40,52]. Briefly, fin clips were digested in 100uL 50mM NaOH at 95⁰C for 30 minutes, and 10uL 1M Tris-HCl pH 8.0 was added after as a buffer. Genotyping occurred through allele-specific forward primers; 5′- CTGGTCATGTGGGTCTCAGC-3 binds to the wild-type allele and 5′- TCTGGTCATGTGGGTCTCATT-3′ binds to the mutant allele. The same reverse primer, 5′- TTTCCAAAGATCACATATCTTGAC-3′, was used for both reactions. The PCR annealing temperature was set to 58⁰C.

### Statistical analysis

As many traits were not normally distributed, non-parametric tests (Mann-Whitney U tests, Kruskal-Wallis tests followed by Dunn’s multiple corrections, and Spearman’s rank correlation tests) were used for conservative estimates of statistical significance. A Hedge’s G test was used to calculate effect size for *oca2* mutant surface fish, as the data had different sample sizes. Analyses were performed and graphs were generated in Graphpad Prism (version 10.3.0, Graphpad Software, graphpad.com).

## Results

### Morphological and behavioral traits differ between surface fish and cavefish from multiple populations

To directly assess the extent to which morphological, behavioral, and physiological traits have repeatedly evolved across cavefish populations, we quantified and compared nine traits between surface fish and cavefish from the three cave populations, Pachón, Molino, and Tinaja, (Fig 1A). First, we assessed bottom-dwelling in response to a novel environment, which is a measure of stress response, and was shown in previous studies to be significantly reduced in cavefish relative to surface fish [27]. Fish from all three cave populations spent significantly less time in the bottom half of the tank compared to surface fish (Fig 1B). Pachón cavefish spent the least amount of time bottom-dwelling, spending significantly less time in the bottom half of the tank than Tinaja cavefish (Fig 1B). Next, we assessed baseline sleep, which is reduced in cavefish [26,47]. Sleep in *A. mexicanus* is defined by periods of inactivity longer than one minute [47]. In agreement with previous studies, we found that sleep is significantly reduced in fish from all cave populations compared to surface fish (Fig 1C).

Cavefish have evolved a number of traits associated with resistance to starvation, including alterations to sleep in response to food deprivation and reduced weight loss following starvation [48,53]. To determine how starvation impacts fish from different cave populations, fish from all populations were starved for 30 days. To assess weight loss and the effects on sleep over this period, we recorded fish weight and sleep prior to and immediately following starvation. While all three cave populations on average lost less weight than surface fish, only Molino cavefish lost significantly less weight than surface fish (Fig. 1D). Fish from all three cave populations show increases in sleep following a period of 30 days starvation, however, only fish from the Tinaja and Molino populations had statistically significant increases in sleep following starvation compared to baseline sleep (Fig S1B-D). In contrast, surface fish did not show a significant directional change in sleep following 30 days of starvation compared to baseline sleep (Fig S1A). Similarly, in all populations, fish that were fed did not show differences in sleep after 30 days compared to baseline sleep levels, confirming there was no effect of retesting (Fig S1). To directly compare starvation-induced changes in sleep between populations, we calculated the change in sleep for each individual starved fish. We found no statistically significant differences between populations in changes in sleep, although change in sleep was increased in fish from Molino and Tinaja populations (Supplemental Fig 2F).

After seven days of recovery from starvation, anesthesia resistance was assessed, as previous work found that cavefish are more resistant to anesthesia than surface fish [39]. When exposed to the anesthetic tricaine, fish from all cave populations exhibited qualitatively increased anesthesia resistance relative to surface fish, with Pachón and Tinaja cavefish taking significantly longer to succumb to anesthesia than surface fish and Molino cavefish (Fig 1E).

Finally, our data demonstrate a shift in morphology consistent with past studies. As reported previously, melanin pigmentation is reduced in Tinaja cavefish and absent in fish from Pachón and Molino populations (data not shown). Cavefish initially develop eyes, which regress over the course of development [25,54]. Fish from all three cave populations have no external eyes or pupils as adults, whereas surface fish have large and fully formed eyes (Fig 1F-G, Fig S2A-B). Cavefish also have expanded numbers of cranial superficial neuromasts compared to surface conspecifics [55–57]. We assessed the number of superficial neuromasts within the bounded area of the suborbital 3 bone. Fish from all three cave populations have significantly more superficial neuromasts than surface fish, and Pachón cavefish have significantly more neuromasts than Molino and Tinaja cavefish (Fig 1H, Fig S2C-E). Together, these data demonstrate that while there are some differences between different cave populations, multiple traits have repeatedly evolved across cavefish populations.

### Evaluation of modes of inheritance in different cave populations

Next, we quantified traits that showed significant differences between surface and cave fish in cave-surface F1 and F2 hybrid fish to determine if modes of inheritance for these traits are shared across cave populations. Similar patterns of inheritance were observed in cave-surface F1 hybrids across all populations for four traits: all F1 hybrids are intermediate between surface and cave phenotypes for eye size (Fig 2E, Fig S3A), pupil size (Fig 2F, Fig S3B), and neuromast number (Fig 2G, Fig S3C-E). Further, all F1 hybrids exhibit statistically significantly reduced anesthesia resistance relative to their corresponding cavefish populations, suggesting a surface-like anesthesia resistance for F1s (Fig 2D). For other traits, however, quantification in surface-cave F1 hybrid fish from different cave populations suggests different inheritance patterns between cave populations. Surface-Pachón and surface-Molino F1 hybrids exhibit surface-like levels of bottom dwelling, while surface-Tinaja F1s exhibit reduced bottom dwelling relative to surface fish, a cave-like phenotype (Fig 2A). Sleep in surface-Pachón F1 hybrids and surface-Molino F1 hybrids is surface-like, while surface-Tinaja F1s appear to be intermediate in phenotype, not significantly different from surface or Tinaja parentals (Fig 2B). We found no statistically significant differences in weight loss between F1 hybrids and either surface or cavefish populations, with the exception of surface-Molino F1 hybrids, which showed a significant increase in weight loss compared to Molino cavefish (Fig 2C). The variance in inheritance across traits in cave-surface F1 hybrids suggests that there is variation between cave populations in the underlying genetic basis of the traits we examined here.

**Figure 2.**
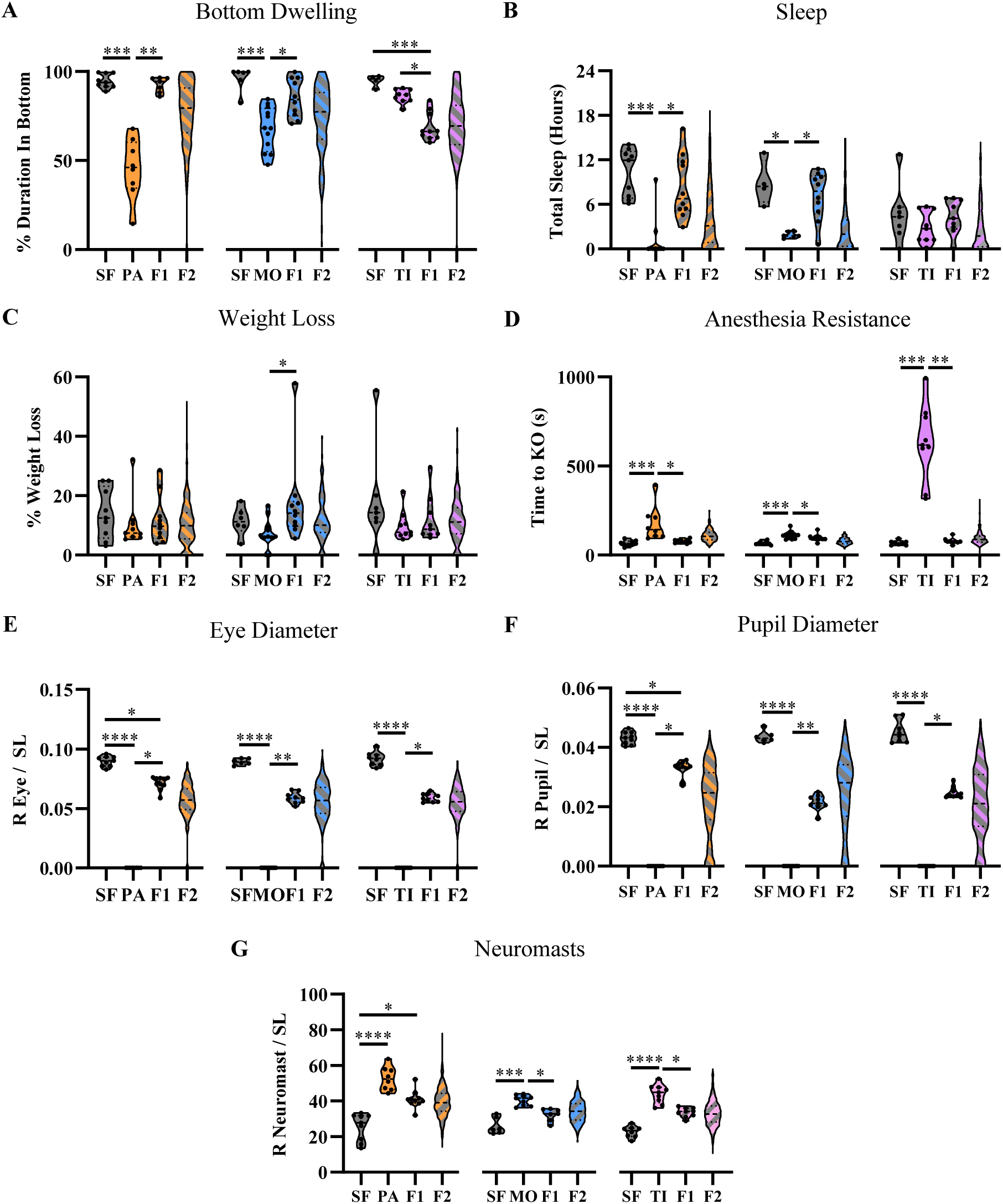
Surface-cave hybrids indicate cave-evolved traits are multigenic. A) Percent time spent bottom-dwelling during the novel tank assay. B) Percent weight lost over 30 days of starvation. C) Time spent awake in the presence of MS-222 anesthetic. D) Dorsal-ventral right eye diameter corrected by standard length. E) Dorsal-ventral right pupil diameter corrected by standard length. F) Number of superficial neuromasts located overtop of the right suborbital 3 bone corrected by standard length. Data points indicate phenotypes of individual fish. Data points were not shown for F2 hybrids. All statistical comparisons were performed between surface fish, cavefish, and F1 hybrids. No statistical comparisons were performed with F2 hybrids. Levels of significance are represented with asterisks p < 0.05 = *, p < 0.01 = **, p < 0.001 = ***, p < 0.0001 = ****.

### Investigating traits in cave-cave hybrids

To further examine whether diverse traits have evolved through similar or different underlying genetic bases, we phenotyped cave-cave F1 hybrids for a subset of traits to assess for complementation. None of the cave-cave hybrids spent more time bottom-dwelling than fish from individual cave populations (Fig S4A). While Pachón-Molino F1 hybrids sleep significantly more than Pachón cavefish, these hybrids spent a similar amount of time sleeping to Molino cavefish, and no increases to more surface-like levels of sleep were observed in cave-cave hybrid fish compared to cavefish from populations from which they derived (Fig S4B). We also did not observe evidence for a shift to reduced anesthesia resistance (Fig S4C), or reduced neuromast number (Fig S4D, Fig S5A-C) in cave-cave hybrids relative to cavefish populations. Similar to previous reports of non-complementation between Molino and Pachón cavefish for albinism [28], our Pachón-Molino hybrid fish were albino (Fig S5D). Both Pachón-Tinaja and Molino-Tinaja F1 hybrids are pigmented, which is consistent with the lack of albinism in Tinaja cavefish and the recessive nature of albinism in both Molino and Pachón populations. However, Pachón-Tinaja and Molino-Tinaja F1 hybrids qualitatively differ in pigmentation (Fig S5D). While eye size was intermediate amongst all three surface-cave F1 hybrids, suggesting additive inheritance, all cave-cave F1 hybrids had no external eyes as adults. Prior research has demonstrated that cave-cave hybrids have larger eyes during larval stages, and have larger, skin-covered internal eyes than either cave parent during adult stages [58–60]. We were unable to quantify eye size in our cave-cave crosses as they can only be measured via euthanization and dissection, since they are not visible externally (Fig S5D). Together, these data suggest that, with the exception of pigmentation in Molino-Tinaja F1 hybrids, no complementation of surface-like traits was observed, suggesting that at least a subset of the same loci may underlie the evolution of these traits across these populations. Together with the patterns of inheritance found in cave-surface crosses, these results suggest both shared and distinct genetic underpinnings contribute to the repeated evolution of traits across these three cavefish populations.

### Genetic correlations are observed across cavefish populations

Next, we investigated the role of shared genetic loci underlying the evolution of multiple traits within cave populations. Analysis of correlations between traits in cave-surface F2 hybrid fish provides a powerful opportunity to investigate whether shared genetic architecture underlies different traits. Previous work in *A. mexicanus* found that multiple traits in the surface-Pachón F2 hybrids covary with one another and have overlapping quantitative trait loci (QTL) peaks in the genome [30,32,33,37]. However, whether cave-derived traits are correlated in other populations of surface-cave hybrids is unknown. To investigate this, we performed pairwise Spearman correlation matrices for traits in each of the three surface-cave F2 hybrid populations. Within each cave population, we identified multiple correlated traits, suggesting that multiple traits evolved together due to linkage or pleiotropy (Fig 3). These correlations were typically weak or moderate, with the exception of eye and pupil size, which were strongly correlated in all three populations (Fig 3), suggesting that shared genetic loci between pairs of traits does not explain all of the genetic architecture underlying these traits within populations.

**Figure 3.**
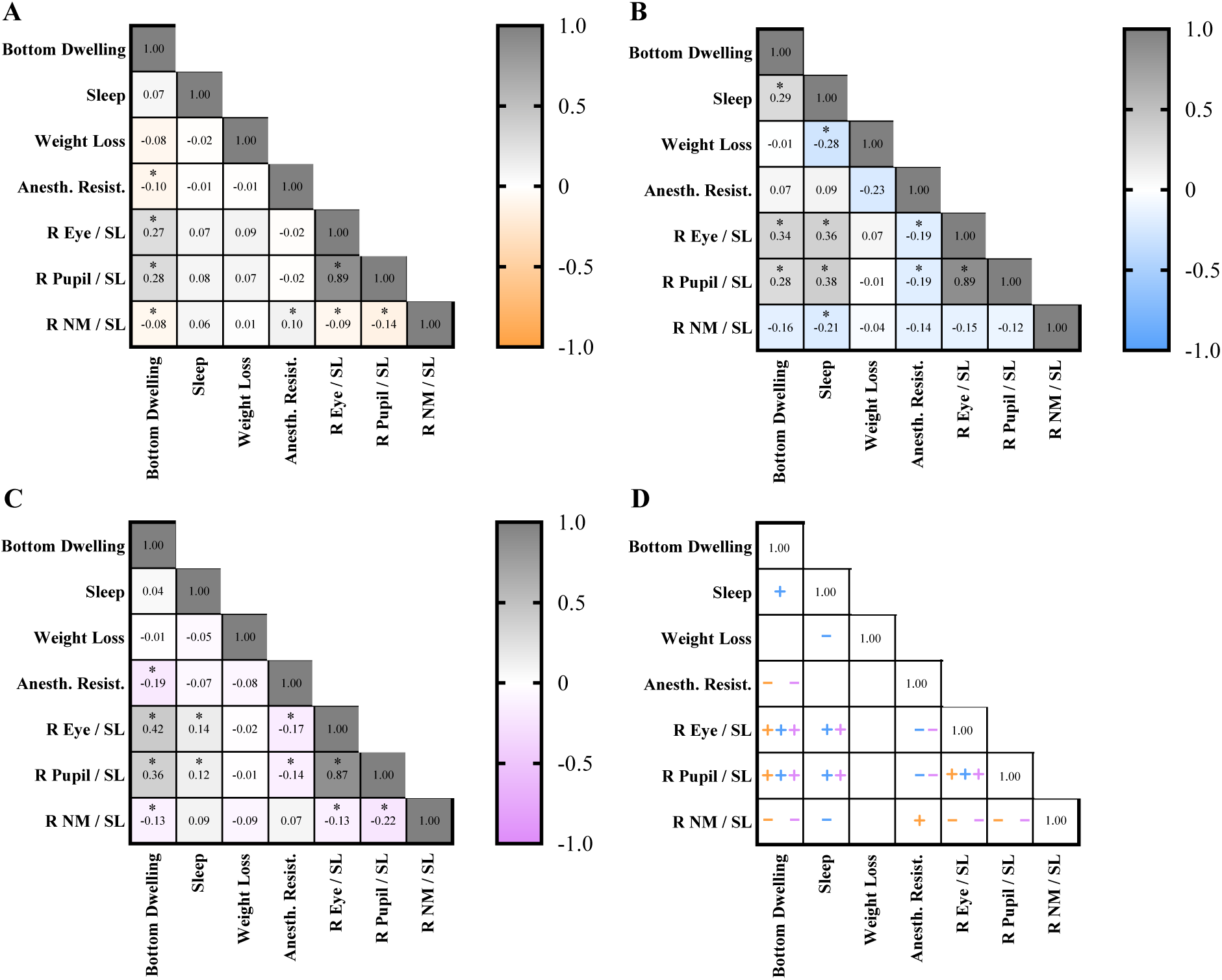
Repeatedly occurring trait covariances in three F2 populations suggest potential shared genetic mechanism for recent trait adaptation. Matrices of Spearman’s rank correlations of traits measured in A) F2 Pachón hybrids, B) F2 Molino hybrids, and C) F2 Tinaja hybrids. Matrices report the Spearman’s r value. Statistically significant covariances are marked with a black internal border. Exact p values and sample sizes are reported on Supplemental Table 1. Levels of significance are represented with asterisks: p < 0.05 = *, p < 0.01 = **, p < 0.001 = ***, p < 0.0001 = ****.

Next, we assessed whether the same traits were correlated across populations. In all three populations, bottom dwelling and eye size, bottom dwelling and pupil size, and eye size and pupil size were all positively correlated (Fig 3). In addition, the directionality of significant correlations for all three cave populations were the same, highlighting the frequency of recurrent covariances (Fig 3D). Multiple correlations were observed between two populations. In surface-Pachón and surface-Tinaja F2 hybrids, bottom dwelling and anesthesia resistance, bottom dwelling and neuromast number, eye size and neuromast number, and pupil size and neuromast number were all negatively correlated (Fig 3A,C). In surface-Molino and surface-Tinaja F2 hybrids, eye size and sleep and pupil size and sleep were positively correlated, and eye size and anesthesia resistance and pupil size and anesthesia resistance were negatively correlated (Fig 3B, C). Several correlations were identified in only one population. In surface-Pachón F2 hybrids, neuromasts and anesthesia resistance were positively correlated (Fig 3A). In surface-Molino F2 hybrids, sleep and bottom dwelling were positively correlated, sleep and neuromast number were negatively correlated, and sleep and weight loss were negatively correlated (Fig 3B).

Finally, we also assessed our surface-Pachón and surface-Molino F2 hybrids for albinism, and investigated whether there were differences in traits between pigmented and albino F2s. Weight loss, anesthesia resistance, and neuromast number were not different between pigmented and albino F2s in either cave population (Fig 4C, D, G). However, albino F2 hybrids derived from both Pachón and Molino populations had smaller pupils and eyes than their pigmented siblings (Fig 4E–F, Supplemental Fig 6A-B). Additionally, albino Molino F2 hybrids exhibited a reduced stress response and sleep relative to pigmented hybrids (Fig 4A-B). Together, these data suggest that shared inheritance between different traits, whether due to linkage or pleiotropy, repeatedly occurs across morphological and behavioral traits.

**Figure 4.**
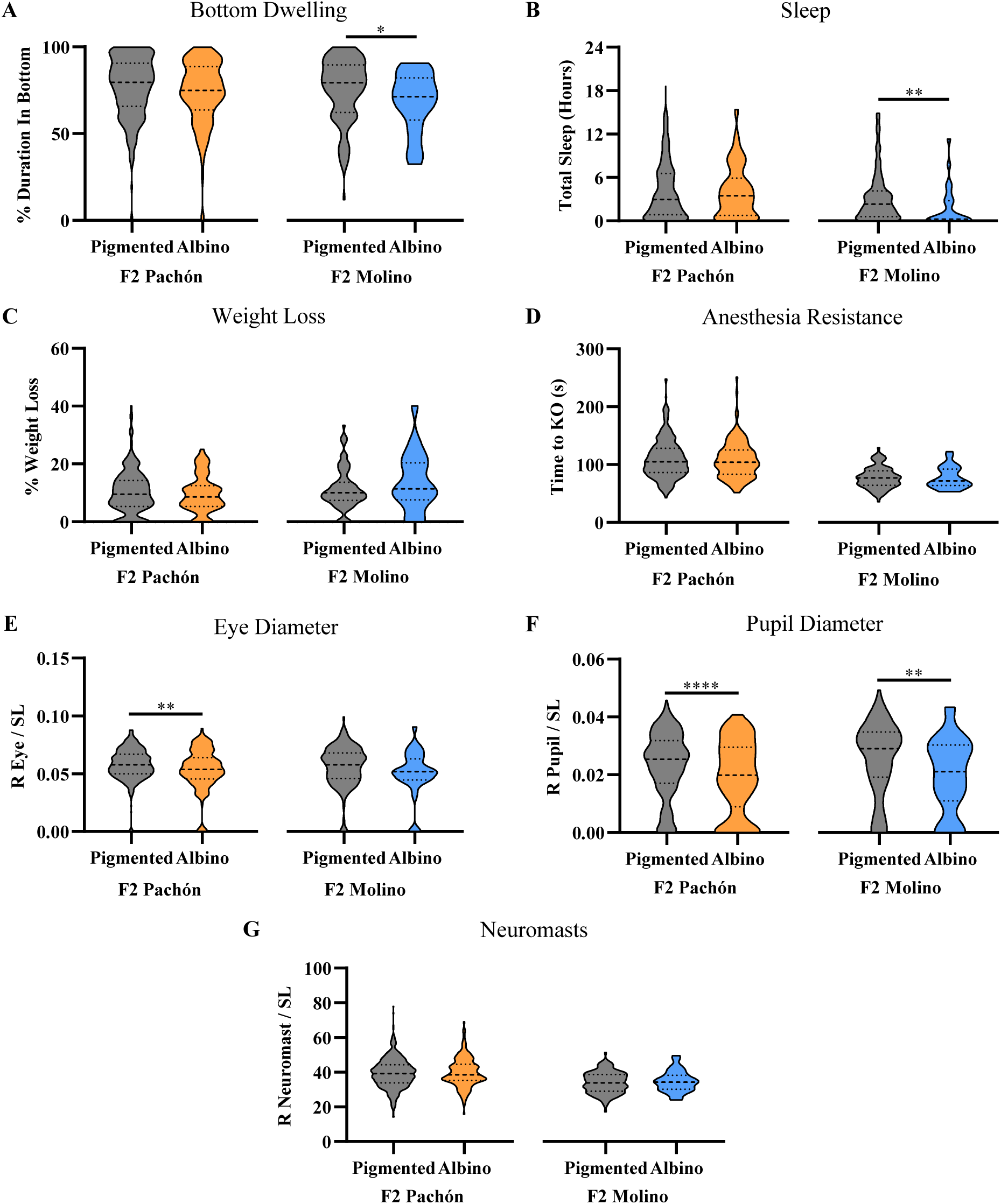
Several traits repeatedly differ between pigmented and albino F2s in Pachón and Molino hybrid populations. A) Percent time spent bottom-dwelling in the novel tank assay. B) Percent weight lost after 30 days of starvation. C) Time spent awake in the presence of tricaine anesthetic. D) Dorsal-ventral right eye diameter corrected by standard length. E) Dorsal-ventral right pupil diameter corrected by standard length. F) Number of superficial neuromasts located overtop of the right suborbital 3 bone corrected by standard length. Levels of significance are represented with asterisks: p < 0.05 = *, p < 0.01 = **, p < 0.001 = ***, p < 0.0001 = ****.

### Directly testing for a role of pleiotropy in observed trait covariances via oca2-mutant surface fish

Traits linked to albinism present a powerful opportunity to directly test whether pleiotropy or linkage impacts evolution of traits. To determine whether *oca2* impacts traits that differed between albino and pigmented hybrids, i.e. bottom dwelling behavior and eye morphology, we utilized a line of surface fish that had CRISPR-Cas9-induced mutations in *oca2* (Fig S7C, [52]). We compared bottom dwelling and eye morphology in wild type (*oca2^+/+^*) and albino (*oca2^Δ2bp/Δ2bp^)* surface fish siblings. Albino surface fish had a reduced bottom dwelling behavior relative to their wild type siblings (Fig 5A) but did not have any significant differences in eye size (Fig 5B, Fig S7A) or pupil size (Fig 5C, Fig S7B), consistent with previous work showing that mutating *oca2* in surface fish larvae is not sufficient to alter eye size [36]. The results from a Hedge’s G test indicate that *oca2* likely has a large effect size on bottom dwelling behavior (G = 0.7484, Table S1). These data suggest that genetic variation at the *oca2* locus may play a role reducing bottom dwelling behavior in these albino cavefish populations.

**Figure 5.**
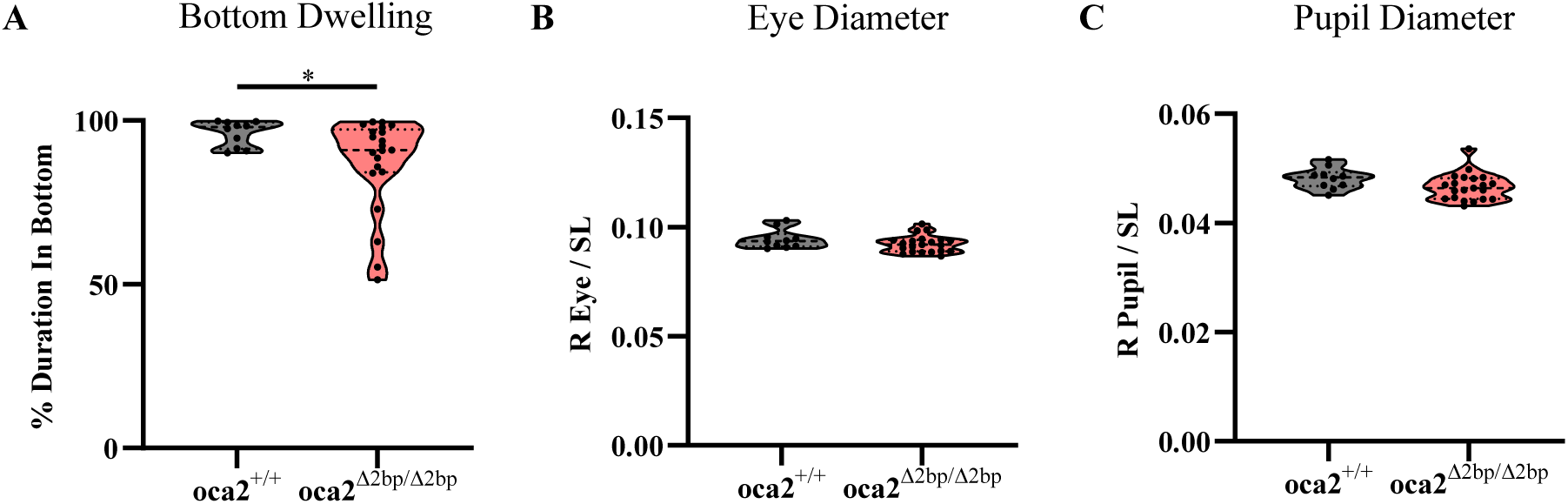
Albino surface fish have smaller pupils than wild type surface fish. A) Percent time spent bottom-dwelling in the novel tank assay between wild type and *oca2^Δ2bp/Δ2bp^* surface fish. B) Dorsal-ventral right eye diameter corrected by standard length between wild type and *oca2^Δ2bp/Δ2bp^*surface fish. C) Dorsal-ventral right pupil diameter corrected by standard length between wild type and *oca2^Δ2bp/Δ2bp^*surface fish. Levels of significance are represented with asterisks p < 0.05 = *, p < 0.01 = **, p < 0.001 = ***, p < 0.0001 = ****.

## Discussion

Repeated phenotypic evolution in response to similar environments provides a powerful opportunity to assess the extent that suites of traits evolve through shared genetic mechanisms both within and between populations. Through quantifying a variety of traits in surface fish, cavefish, hybrid crosses, and *oca2* mutant surface fish, we demonstrate that many cave-derived traits are correlated, suggesting that shared genetic architecture, via pleiotropy or close linkage, has repeatedly contributed to the evolution of the cave phenotype.

### Shared inheritance and concentrated genetic architectures

Suites of traits often evolve together when populations experience strong selection, as selection acts on combinations of traits that jointly influence fitness rather than on isolated phenotypes. Cave animals across taxa have repeatedly evolved suites of similar traits, including reductions in the visual system, loss or reduction of pigmentation, and enhancement of non-visual sensory systems [61]. Similarly, multiple morphological, physiological and behavioral traits have evolved in *A. mexicanus* cavefish populations [29,62]. Thus, cave animals provide a powerful system in which to study multi-trait evolution. Across multiple independent cavefish populations, we identified correlations among traits such as eye and pupil size, stress response, neuromast expansion, and anesthesia resistance in cave-surface F2 hybrids, consistent with a role for concentrated genetic architectures underlying multiple traits involved in cave adaptation. Our results extend earlier work showing correlations for distinct traits in Pachón hybrids [30,31,33]. These prior works also identified overlapping QTL for their traits, and future QTL mapping of our dataset will give insight into the commonality of overlapping QTL in populations outside Pachón.

### Pleiotropy, linkage, and the case study of oca2

Determining if co-variation between traits is due to pleiotropy can be challenging, as differentiating between pleiotropy and close linkage is difficult [63]. Indeed, multi-trait evolution through concentrated genetic loci has been attributed to pleiotropy and to linkage in other species. For example, a repeatedly identified mechanism underlying multi-trait evolution is evolution of traits via supergenes, chromosomal inversions that contain multiple genes in which multiple individual genes separately impact different traits [17]. Supergenes have been associated with multi-trait evolution across species, including redpoll finches, butterflies, and deer mice [64–66]. Conversely, multiple instances of pleiotropic genes contributing to the evolution of traits have been identified. One of these is the *foraging* gene in *Drosophila melanogaster,* which was initially identified as playing a role in larval foraging behavior, and has since been implicated in natural variation numerous other traits, including learning and memory, sleep and temperature tolerance, among others [67].

The genes and genetic changes that contribute to the evolution of traits in cave *A. mexicanus* are largely unknown, however, the basis for albinism is known to occur through *oca2* [28,52]. This provides the opportunity to investigate traits covarying with albinism to differentiate between pleiotropy and close linkage. Here, we performed a direct test of the role of *oca2* in multi-trait evolution. In addition to the role *oca2* plays in albinism in cavefish [28,52,68], *oca2* has been implicated in catecholamine levels, anesthesia resistance, larval feeding behavior, and sleep loss [36,38–40]. We found that albino hybrids differed in stress response, sleep, and eye/pupil size relative to pigmented siblings. Examination of two of these traits in *oca2* mutant surface fish confirmed that *oca2* impacts bottom-dwelling, but not eye/pupil size. This suggests a mixture of true pleiotropy (direct effects of *oca2* on stress behavior) and linkage (eye size influenced by loci nearby), however, we cannot rule out the possibility that while not sufficient to alter eye size when mutated alone in surface fish, *oca2,* in combination with other genetic changes, impacts eye and/or pupil size in cavefish. The coexistence of these mechanisms aligns with theoretical predictions that adaptation can be facilitated by both pleiotropy and physical clustering of alleles [8]. Whether pleiotropy or linkage play a role in the co-variation of other cave phenotypes identified here remains to be seen.

### Repeated evolution and alternative evolutionary paths

Across all three cave-surface F2 hybrid populations, we find both shared and population-specific correlations among traits, suggesting that some aspects of cave evolution are shaped by common genetic architectures while others have evolved through distinct, lineage-specific correlations. Across all three populations, we found correlations between eye phenotypes and bottom dwelling, similar to the direction of trait change seen in cavefish evolution. Several trait correlations were present only in Pachón and Tinaja, and several additional trait correlations were observed between Molino and Tinaja as well. We also observed population-specific correlations, particularly in Molino hybrids. These results suggest that while some multi-trait correlations are common across populations, others have evolved only in specific cave populations, indicating that both shared and lineage-specific genetic architectures underlie repeated evolution. Whether the same or distinct loci underlie trait correlations observed across different cave populations remains an open question. Both crosses and genetic mapping studies suggest that for many traits in cavefish, both shared and distinct genetic loci contribute to the repeated evolution of these traits [28,34,60]. Genetic mapping studies of these traits in multiple cave populations may provide additional insight into whether repeated instances of co-variation across cave populations are due to the same or different genomic regions.

Notably, the correlations amongst traits we observe align with the directions of phenotypic evolution that we repeatedly observe in cavefish, suggesting that the genetic variance–covariance matrix is at least partly structured along axes that facilitate the direction of phenotypic evolution we observe in the caves [69]. One caveat is that while phenotypic correlations in F2s provide insight into the potential shared genetic bases, shared environmental conditions within the laboratory environment may influence the phenotypic correlations we observe here.

In conclusion, our findings suggest that in *A. mexicanus,* concentrated genetic architectures, due to pleiotropy or close linkage, underlie multiple cave-adapted traits. Moreover, we find that multi-trait evolution via these concentrated genetic architectures occurs across different cave populations. Through functional validation, we find that *oca2* influences bottom dwelling through pleiotropy and is likely closely linked with another loci that influences eye size and pupil size.

Future studies that integrate QTL mapping and further functional validation can help identify cases of pleiotropy versus linkage and shed additional light on the genetic basis of repeated evolution.

## Acknowledgements

This work was supported by National Institutes of Health awards R35GM138345 to JEK and R24OD030214 to ACK and NSF IOS 1933076/2202359 to SEM, JEK.

