## Supplemental Figures for "Analysis of multi-trait evolution across independently evolved cavefish populations reveals shared and independent evolution of suites of cave-associated traits"

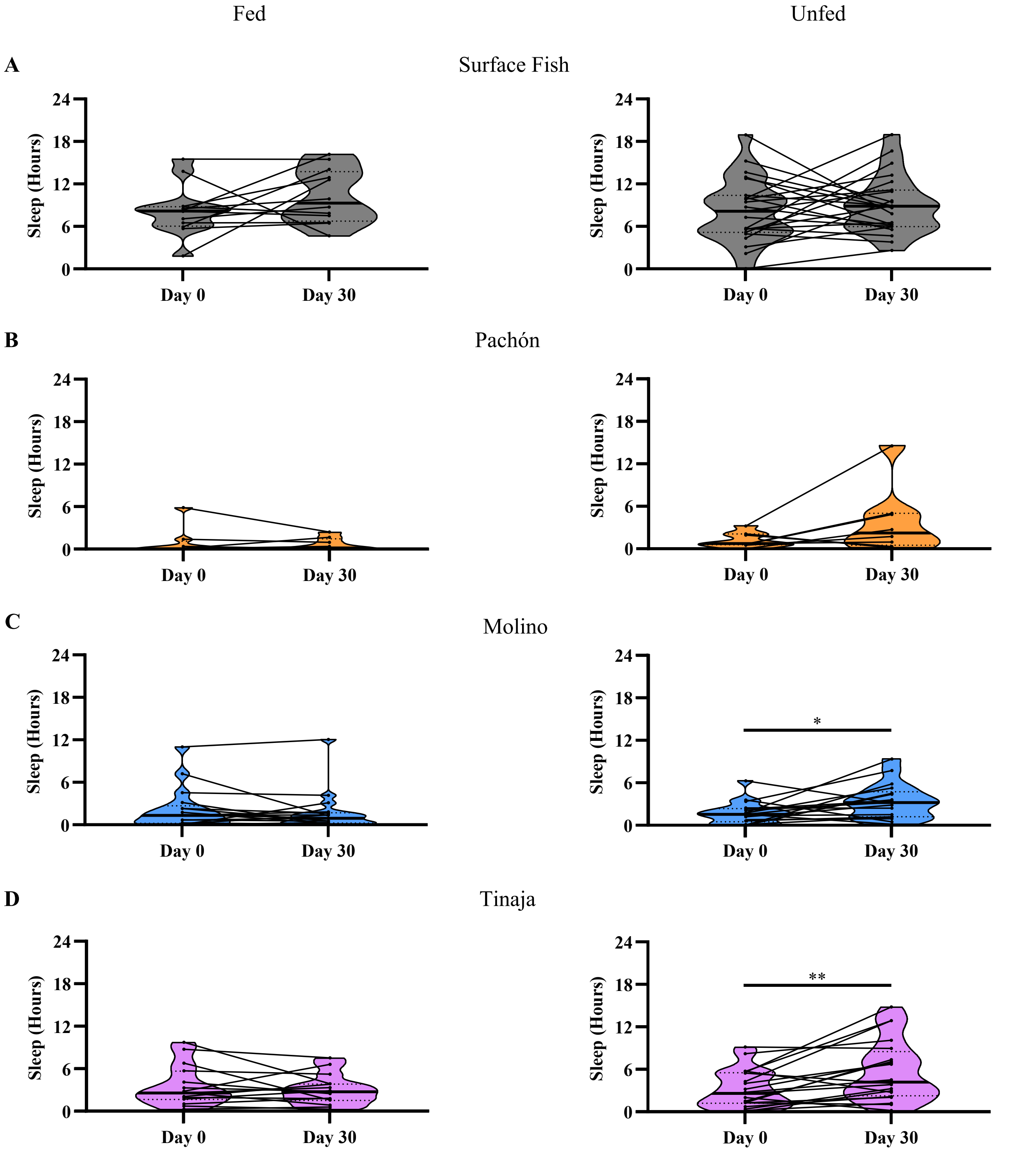


**Supplemental Figure 1. Change in sleep over 30 days in Surface, Pachón, Molino, and Tinaja populations.** A) Change in total sleep in fed and unfed surface fish. B) Change in sleep in fed and unfed Pachón fish. C) Change in sleep in fed and unfed Molino fish. D) Change in sleep in fed and unfed Tinaja fish. Levels of significance are represented with asterisks: $p < 0.05$ = *, $p < 0.01$ = **, $p < 0.001$ = ***, $p < 0.0001$ = ****.

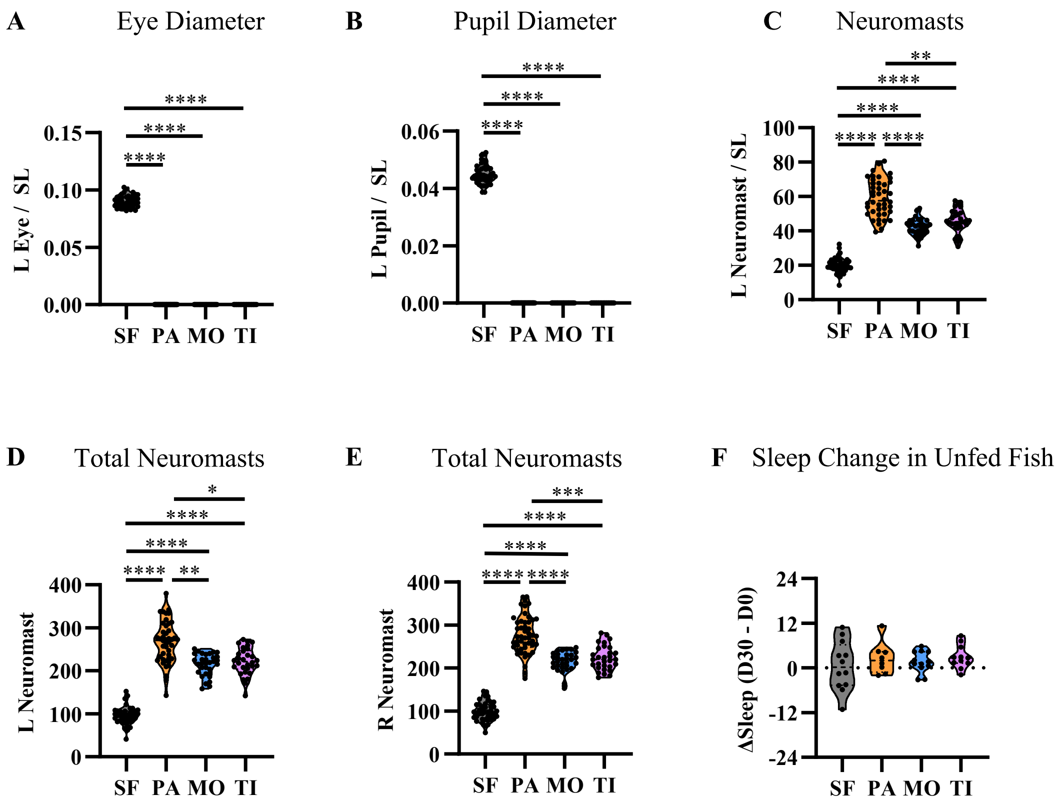


**Supplemental Figure 2. Left-sided morphological data across *A. mexicanus* populations.** A) Dorsal-ventral measurements of the left eye diameter, corrected by standard length. B) Dorsal-ventral measurements of the left pupil diameter, corrected by standard length. C) Number of superficial neuromasts overtop of the left suborbital 3 bone, corrected by standard length. D) Number of superficial neuromasts overtop of the left suborbital bone, uncorrected by standard length. E) Number of superficial neuromasts overtop of the right suborbital bone, uncorrected by standard length. F) Change in sleep over 30 days of starvation in surface, Pachón, Molino, and Tinaja fish. Levels of significance are represented with asterisks: p < 0.05 = *, p < 0.01 = **, p < 0.001 = ***, p < 0.0001 = ****.

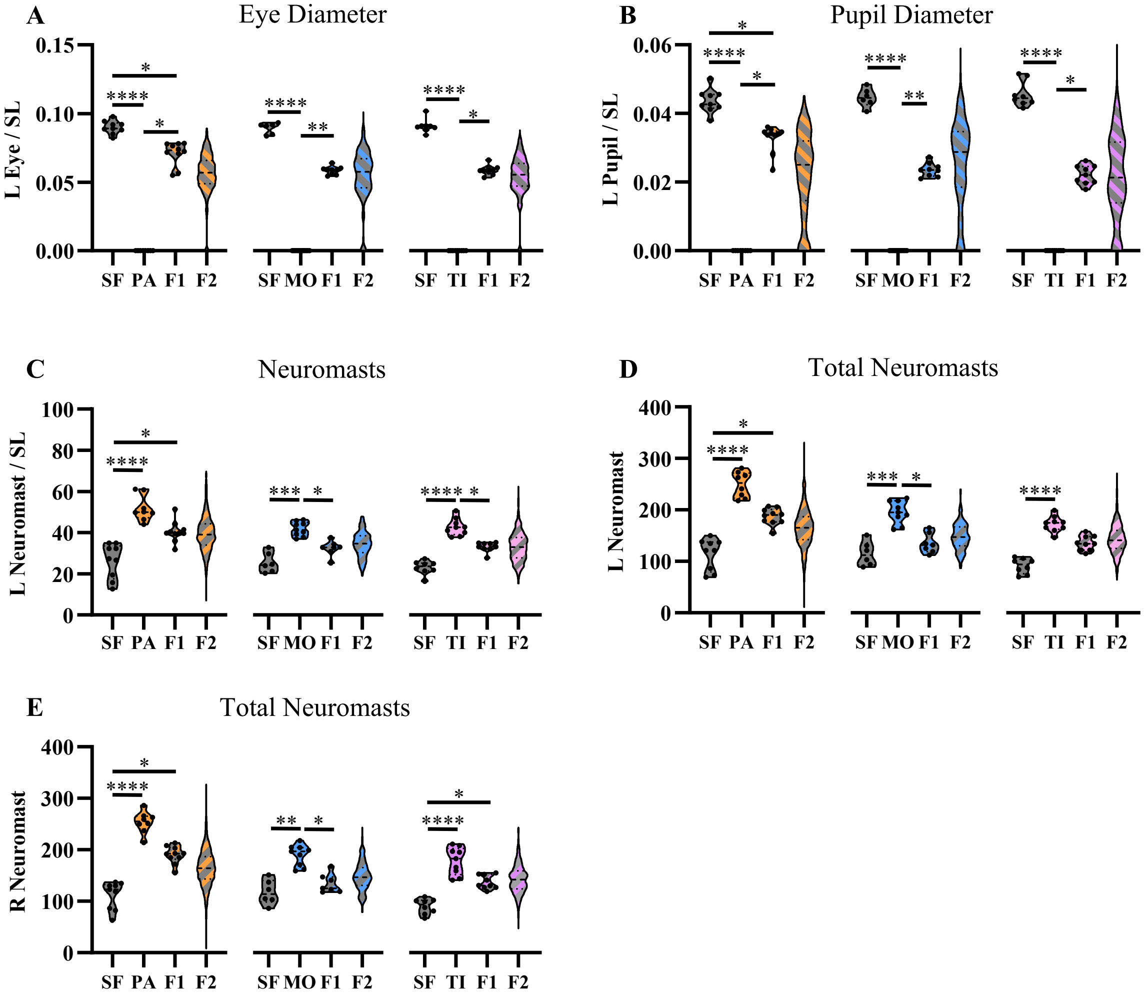


**Supplemental Figure 3. Left-sided morphological data in parental, F1 hybrid, and F2 hybrid populations.** A) Dorsal-ventral measurements of the left eye diameter, corrected by standard length. B) Dorsal-ventral measurements of the left pupil diameter, corrected by standard length. C) Number of superficial neuromasts overtop of the left suborbital 3 bone, corrected by standard length. D) Number of superficial neuromasts overtop of the left suborbital bone, uncorrected by standard length. E) Number of superficial neuromasts overtop of the right suborbital bone, uncorrected by standard length. Statistics were performed only on surface, cave, and F1 hybrid populations. Levels of significance are represented with asterisks: $p < 0.05$ = *, $p < 0.01$ = **, $p < 0.001$ = ***, $p < 0.0001$ = ****.

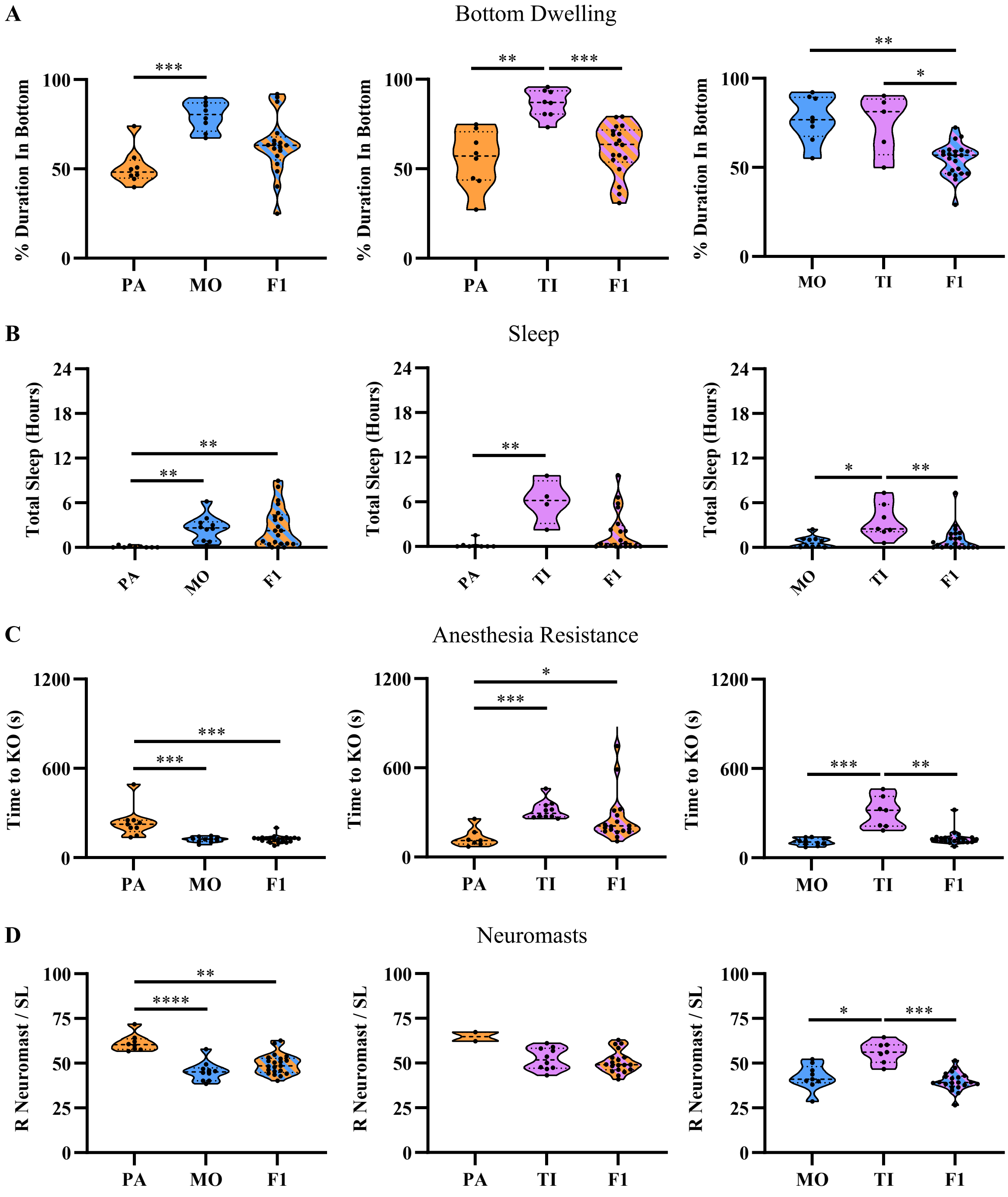


**Supplemental Figure 4. Traits in cave and cave-cave F1 hybrid populations.** A) Percent duration spent in bottom of tank during the novel tank assay in cave and F1 hybrid populations. B) Total sleep duration over 24 hours in cave and F1 hybrid populations. C) Time spent awake in the presence of MS-222 anesthetic in cave and F1 hybrid populations. D) Number of superficial neuromasts overtop of the right suborbital 3 bone, corrected by standard length. Levels of significance are represented with asterisks: $p < 0.05$ = *, $p < 0.01$ = **, $p < 0.001$ = ***, $p < 0.0001$ = ****.

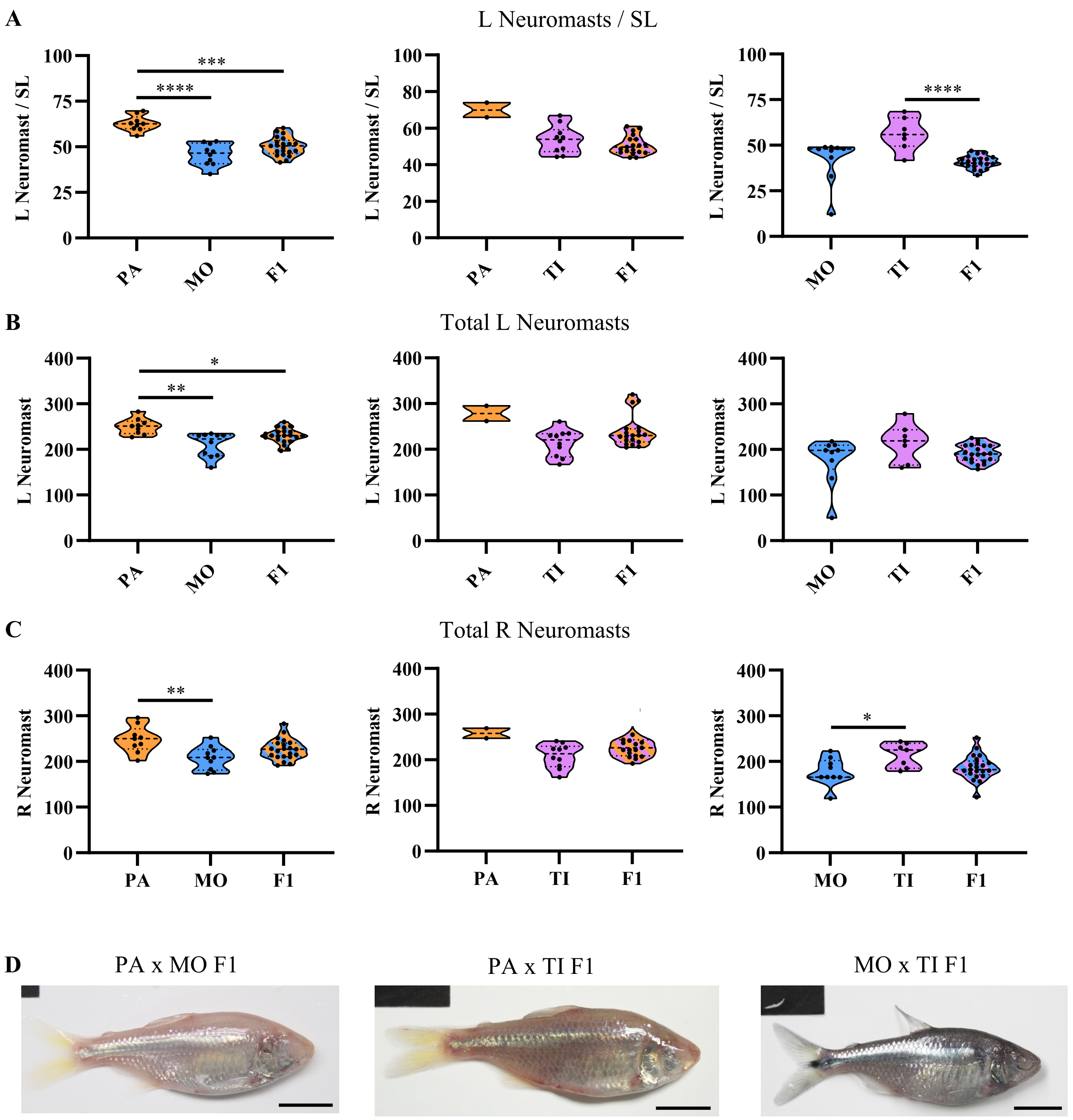


**Supplemental Figure 5. Left-sided neuromasts in cave and cave-cave F1 hybrid populations.** A) Number of superficial neuromasts overtop of the left suborbital 3 bone, corrected by standard length. B) Number of superficial neuromasts overtop of the left suborbital 3 bone, uncorrected by standard length. C) Number of superficial neuromasts overtop of the right suborbital 3 bone, uncorrected by standard length. D) Full-body images of Pa x MO F1s, PA x TI F1s, and MO x TI F1s. Levels of significance are represented with asterisks: $p < 0.05$ = *, $p < 0.01$ = **, $p < 0.001$ = ***, $p < 0.0001$ = ****.

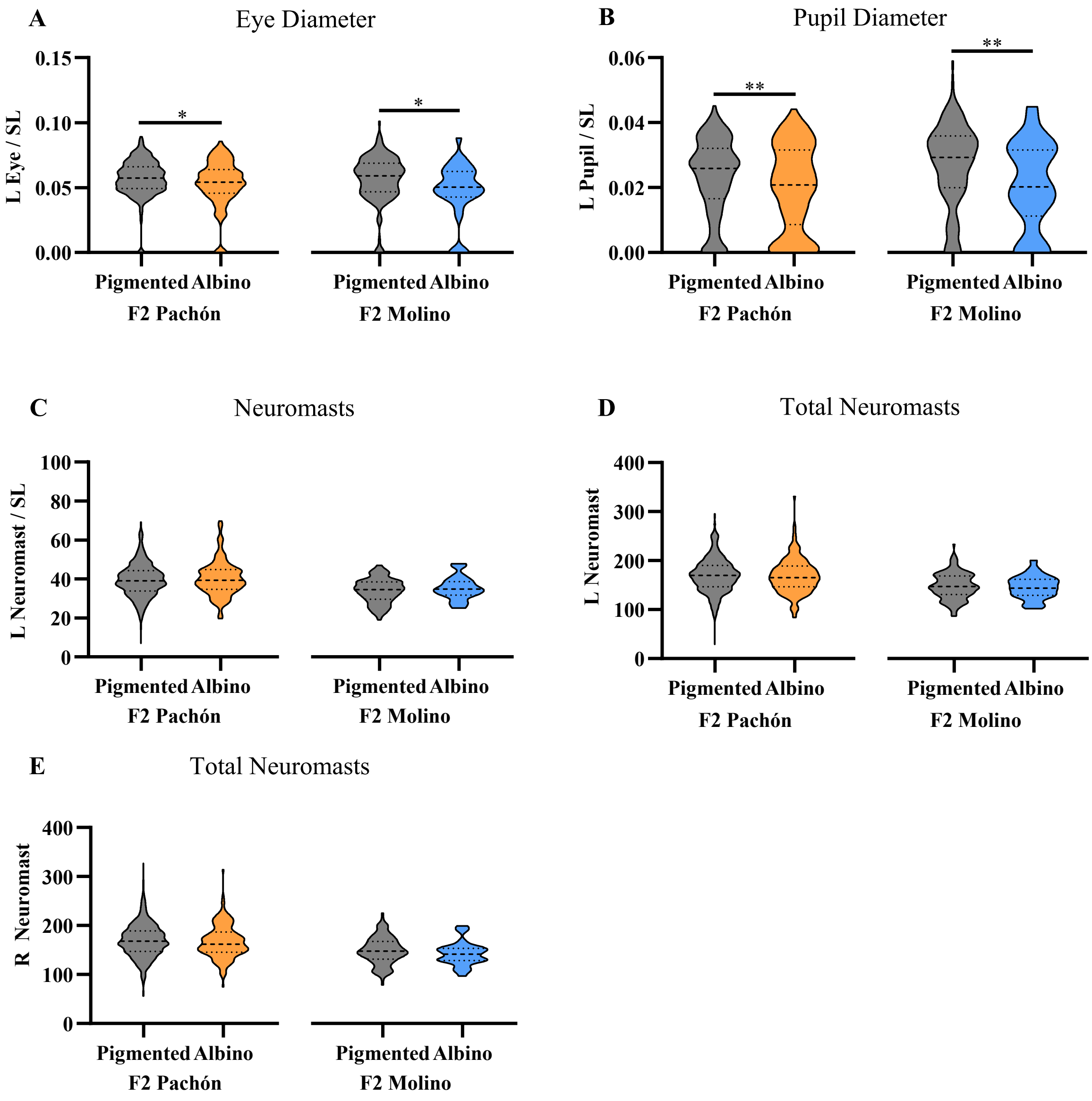


**Supplemental Figure 6. Left-sided morphological traits in F2 Pachón and F2 Molino hybrids, split by pigmentation.** A) Dorsal-ventral measurements of left eye diameter, corrected by standard length. B) Dorsal-ventral measurements of the left pupil diameter, corrected by standard length. C) Number of superficial neuromasts overtop of the left suborbital 3 bone, corrected by standard length. D) Number of superficial neuromasts overtop of the left suborbital 3 bone, uncorrected by standard length. E) Number of superficial neuromasts overtop of the right suborbital 3 bone, uncorrected by standard length. Levels of significance are represented with asterisks: $p < 0.05$ = *, $p < 0.01$ = **, $p < 0.001$ = ***, $p < 0.0001$ = ****.

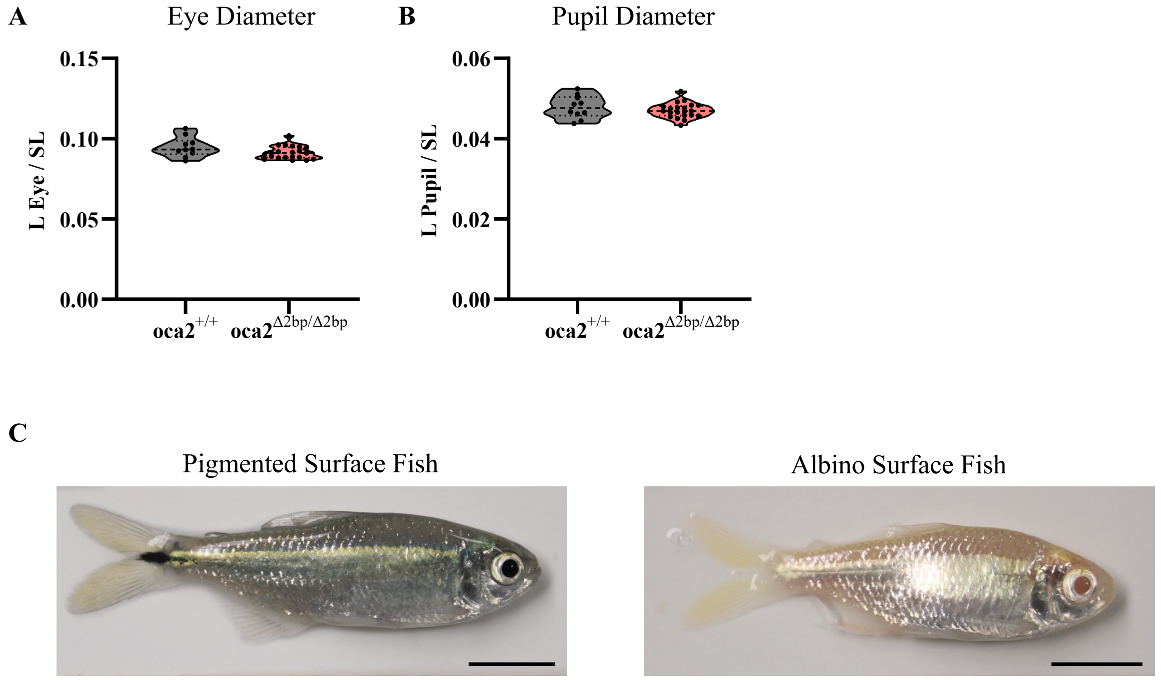


**Supplemental Figure 7. Left-sided eye morphology in wild type and *oca2* mutant surface fish.** A) Dorsal-ventral measurements of left eye diameter, corrected by standard length. B) Dorsal-ventral measurements of the left pupil diameter, corrected by standard length. C) Full-body images of pigmented and albino surface fish siblings. Levels of significance are represented with asterisks: $p < 0.05$ = *, $p < 0.01$ = **, $p < 0.001$ = ***, $p < 0.0001$ = ****.
